# Type 2 Diabetes Mellitus pathogenesis and role of Peptidylglycine Alpha-Amidating Monooxygenase (PAM) Gene elaboration by In-silico Analysis

**DOI:** 10.1101/2025.10.12.681891

**Authors:** Muhammad Mohsin, Harliana Mohd Yusof

**Affiliations:** MSc Medical Bioscience, School of Health and Life Sciences, Glasgow Caledonian University, Glasgow, UK; Oncology Department, Portsmouth Hospitals University NHS Trust, Portsmouth, UK

**Keywords:** PAM gene, Type 2 Diabetes Mellitus, in-silico, bioinformatics, IUPred-3, InterPro, dbSNP, CRISPR

## Abstract

Type 2 Diabetes Mellitus (T2DM) is caused by pancreatic beta cell failure and alpha cell dysfunction that are central to this disease pathophysiology. For T2DM, the Peptidylglycine Alpha-Amidating Monooxygenase (PAM) gene has susceptible locus as identified by Genome-wide association studies (GWAS) though underlying molecular mechanisms remained poorly characterised. This study have explored comprehensive in-silico characterization of the PAM gene potential role in T2DM. InterPro was employed to identify protein family and it was observed it belongs to Peptidylglycine alpha-hydroxylating monooxygenase/peptidyl-hydroxyglycine alpha-amidating lyase family (IR000720), consisting of two catalytic domains and nine active regions. Clinically significant 153 genetic variants including non-synonymous SNPs within the locus were identified after analysis of dbSNP through UCSC Genome Browser. Intrinsically disrupted large region (aa 290-495) was observed during protein disorder prediction by utilizing IUPred-3. Ensembl and NCBI BLAST were used to analyse evolutionary conservation sites. This analysis resulted in high sequence similarity with common model organisms Mus musculus and Rattus norvegicus with similarity 90% and 89% respectively. These computational findings suggest specific PAM variants likely to disrupt protein function, providing validated future direction for experimental studies to confirm PAM gene pivotal role in T2DM.

## Introduction

Pancreatic beta cells failure due to insensitivity of insulin due to insulin resistance which results in decreased insulin manufacturing. This render into abnormal glucose transportation towards the different body organs and tissues like liver, muscle cells and adipose tissues. In recent studies, it has been proven that alpha-cells also play an important role in the pathophysiology of the T2DM.

In T2DM, glucose and glucagon levels that incline during fasting, are not inhibited due to meal intake. Hyperglycemia results due to the increased insulin resistance and not enough levels of the insulin. Occurrence of type 2 diabetes mellitus (T2DM) as chronic metabolic syndrome is growing gradually worldwide. It is being converted into epidemic disease in some countries. It is due to people suffering from T2DM are anticipated to be two-fold in coming decade due to ageing of people will increase (Olokoba et al., 2012). Last few decades of 20th century had shown T2DM is clear and persistent problem for developed and for developing world (Ginter and Simko, 2013).

Patients suffering from the T2DM may get concurrent diseased conditions like microvascular and macrovascular complications. Neuropathy, nephropathy and retinopathy are the microvascular anomalies. Metabolic syndrome and cardiovascular conditions are macrovascular comorbidities. T2DM can be induced due to environmental factors like unhealthy diet, lack of exercise and obesity. Glucose homeostasis is influenced by the pathophysiological changes due to genetic factors(DeFronzo et al., 2015).

There is evident association of inherited genetic abnormalities with first degree type 2 diabetes mellitus. Inheritable genetic anomalies cause T2DM significantly. Monozygotic twins have almost 100 percent concordance and about 25 percent get T2DM who have T2DM in their family history(Olokoba et al., 2012)

Peptidylglycine Alpha-amidating Monooxygenase (PAM)

Biosynthesis of many endocrine and neural peptides is facilitated by this bifunctional enzyme that play role in modification of non-active peptidylglycine precursors into the related active form of alpha-amidated peptide through post translational catalysis (Satani et al., 2003).

## Methods

### Gene and Protein Identification

NM_000919.4 canonical PAM transcript served as reference sequence to perform all analyses. Ensembl genome database (Release 104) was used to locate gene on chromosome 5q21.1, catalogue its isoforms, and assessed its evolutionary conversations by orthologue mapping.

### Protein Domain and Family Analysis

Using the InterPro database (Blum et al., 2021), protein family membership was determined along with identifying the functional domains, conserved active sites were indicated within the PAM amino acid sequence.

### Genetic Variation Analysis

NCBI dbSNP database (Sherry et al., 2001) was used to retrieve genetics variants that included single nucleotide polymorphisms (SNPs). UCSC Genome Browser (Kent et al., 2002) was deployed on the Human Dec. 2013 (GRCh38/hg38) assembly to visualize and analyse the genetic variants. SNPs presented in database, as clinically flagged genetic variants, were explicitly observed.

### Sequence and Structural Analysis

UniprotKB(accession P19021) (UniProt Consortium, 2023) tool was used to obtain canonical amino acid sequence for the PAM gene protein. Afterward, fundamentally disorder protein regions within its structure were predicated by using the IUPred-3 web server (Erdős and Dosztányi, 2021). Modbase (Pieper et al., 2014) was used to visualized three dimensional protein structure.

### Evolutionary Conservation

Comparative sequence analysis was carried out to evaluate evolutionary conservation between the human PAM protein and its orthologues in model organisms (Mus musculus and Rattus norvegicus) by NCBI BLAST tool (Johnson et al., 2008; Altschul et al., 1990).

### Hypothetical Experimental Design

CHOPCHOP web tool (Labun et al., 2019) was used to design guide RNA(sgRNA) sequence for “CRISPR-Cas9-mediated” PAM gene editing in PAM locus in mouse model.

## Results

### Protein Domain Architecture and Functional Regions

Analysis with InterPro revealed that the PAM gene belongs to the Peptidylglycine alpha-hydroxylating monooxygenase/peptidyl-hydroxyglycine alpha-amidating lyase family (IR000720). The protein contains two essential catalytic domains - Peptidylglycine alpha-hydroxylating monooxygenase (PHM) and Peptidylglycine amidoglycolate lyase (PAL) - which work sequentially in the peptide amidation process. Nine distinct active regions were identified within the amino acid sequence at positions: 18-36, 91-112, 132-151, 255-274, 301-320, 523-541, 587-606, 629-652, and 783-806.

**Figure 1.**
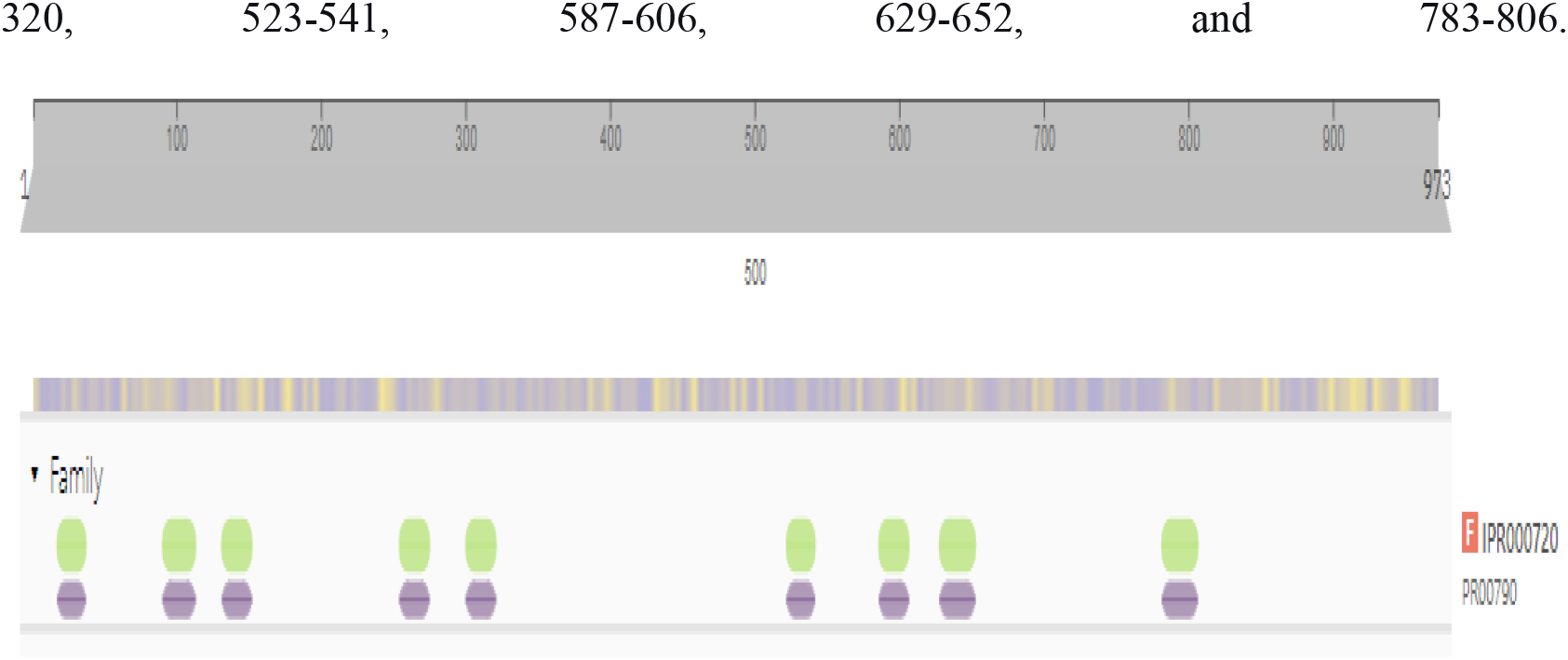
InterPro analysis showing PAM protein family classification and active regions.

**Figure 2.**
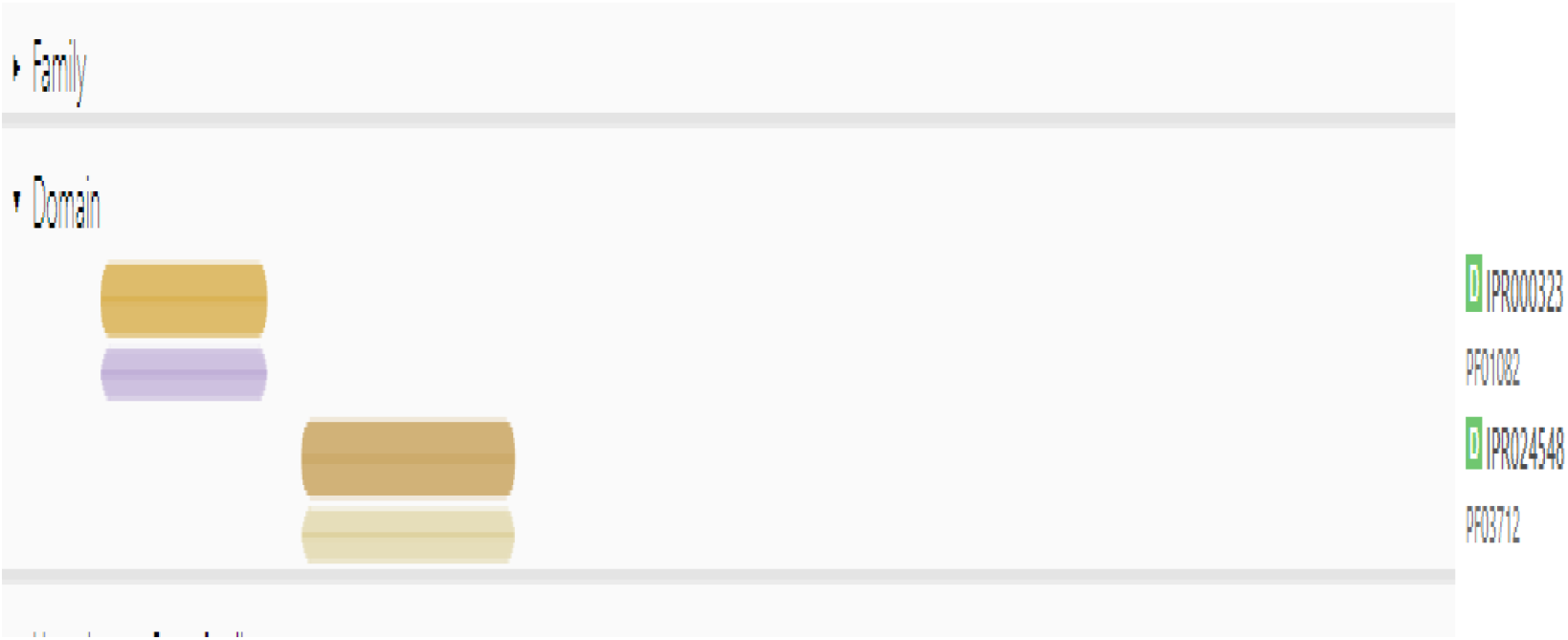
Domain architecture of PAM protein showing catalytic domains.

**Figure 3.**
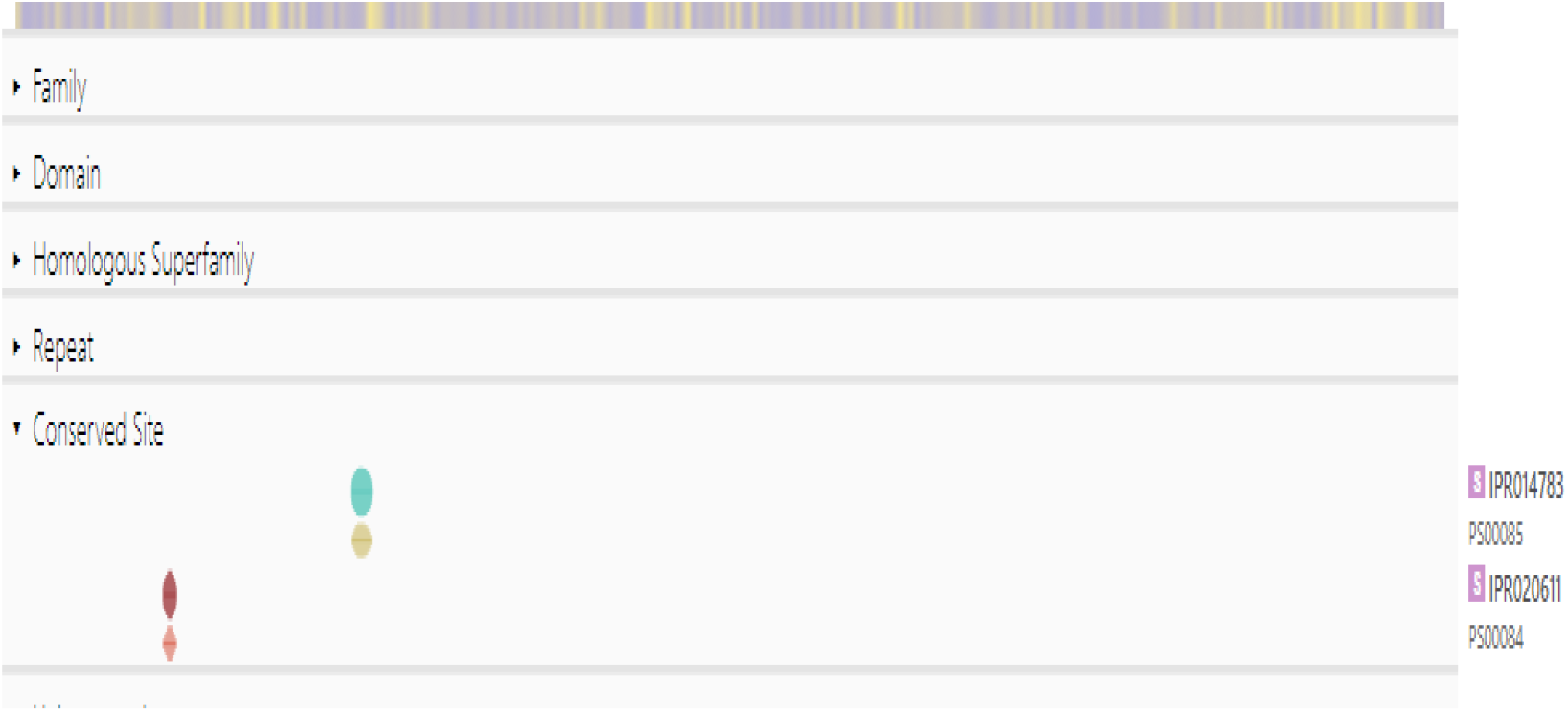
Conserved metal-binding sites in PAM protein.

### Genetic Variant Analysis

Comprehensive analysis of dbSNP through UCSC Genome Browser identified 153 genetic variants within the PAM locus. Among these, 142 were simple nucleotide polymorphisms, including several non-synonymous SNPs that result in amino acid changes. Notably, multiple variants were clinically flagged by NCBI as potentially significant, with several located within critical functional domains that could impact enzymatic activity.

**Figure 4.**
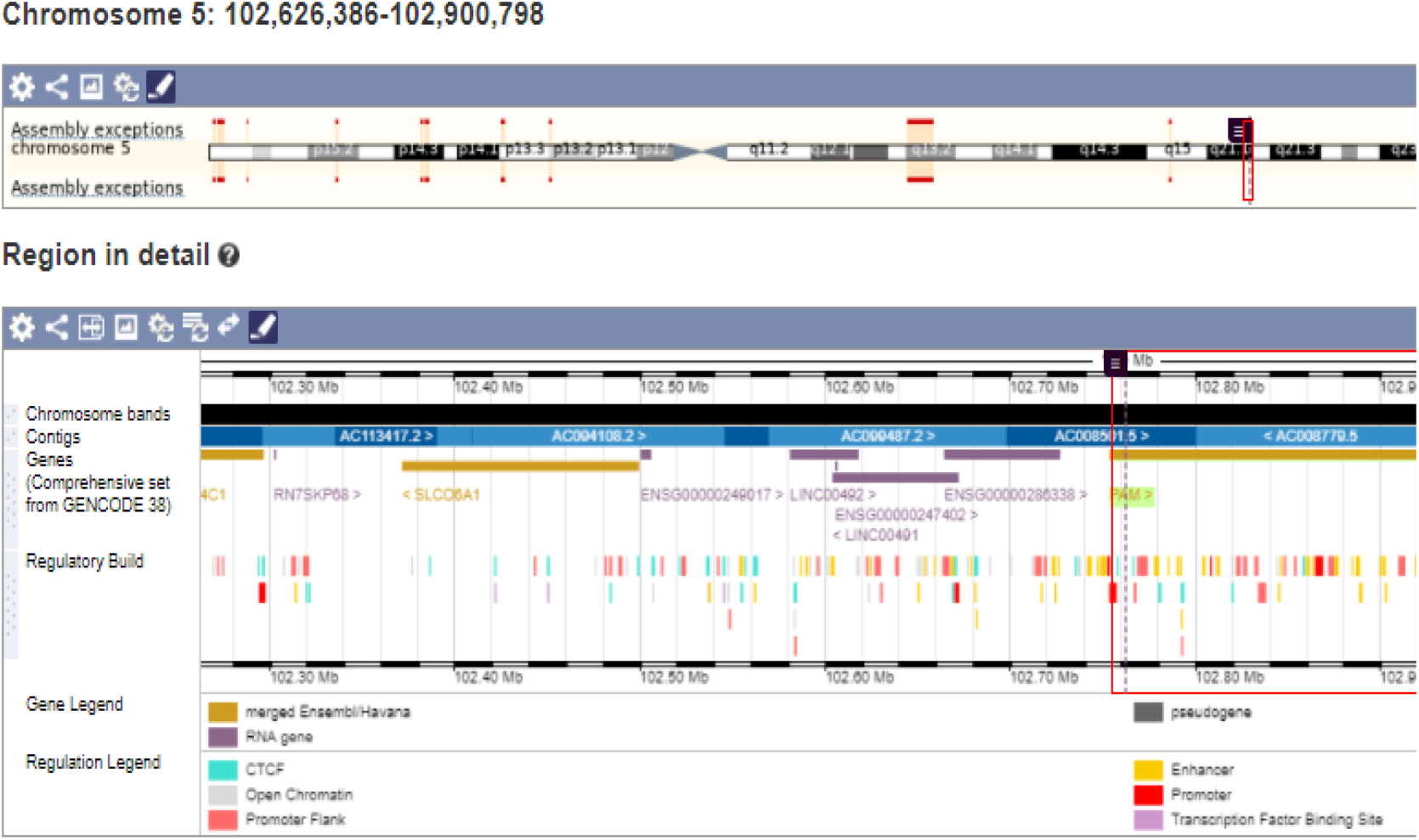
UCSC Genome Browser view showing PAM gene location on chromosome 5.

**Figure 5.**
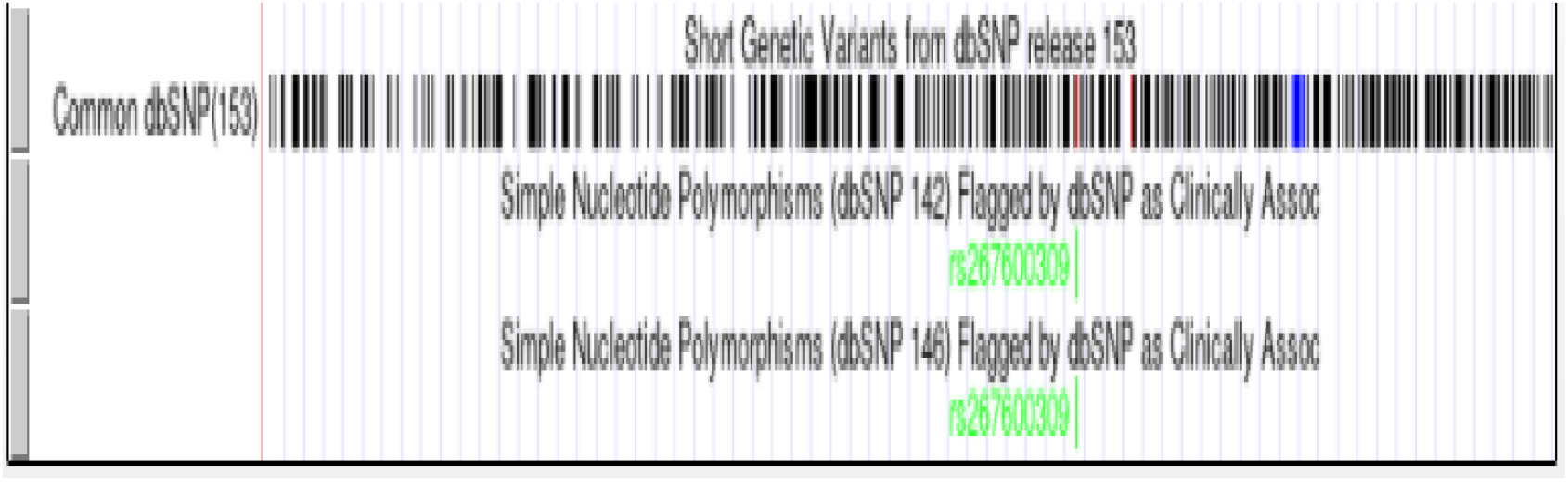
dbSNP variants mapped to PAM gene structure.

**Figure 6.**
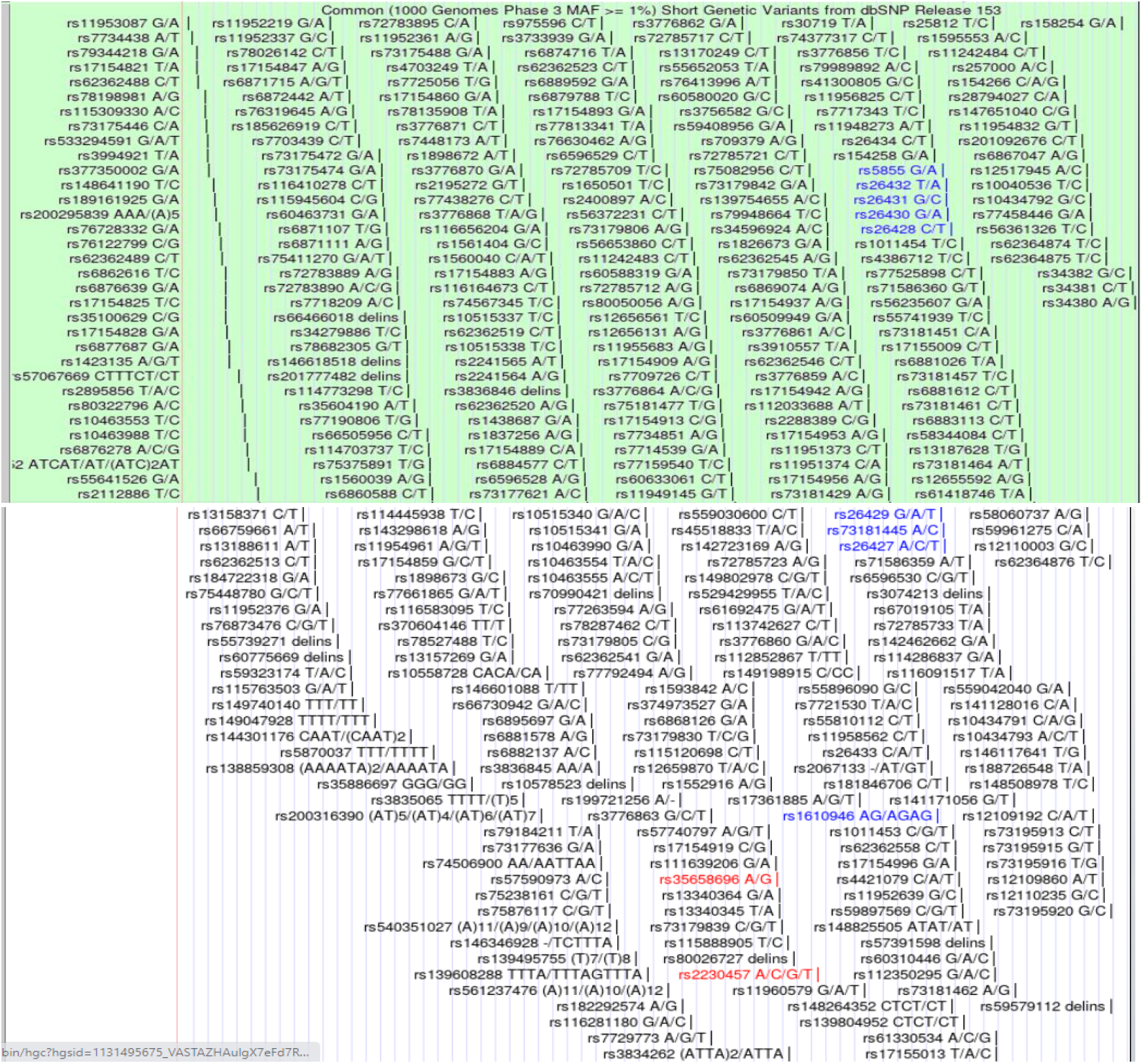
Detailed view of translated regions and amino acid changes.

### Protein Structure and Disorder Prediction

Modbase analysis provided the three-dimensional structure of PAM protein, revealing key structural features relevant to its enzymatic function. IUPred-3 analysis predicted a large intrinsically disordered region spanning amino acids 290-495, with disorder scores consistently above 0.5. This extensive disordered region suggests potential roles in protein-protein interactions and structural flexibility that may be important for PAM’s function in secretory pathways.

**Figure 7.**
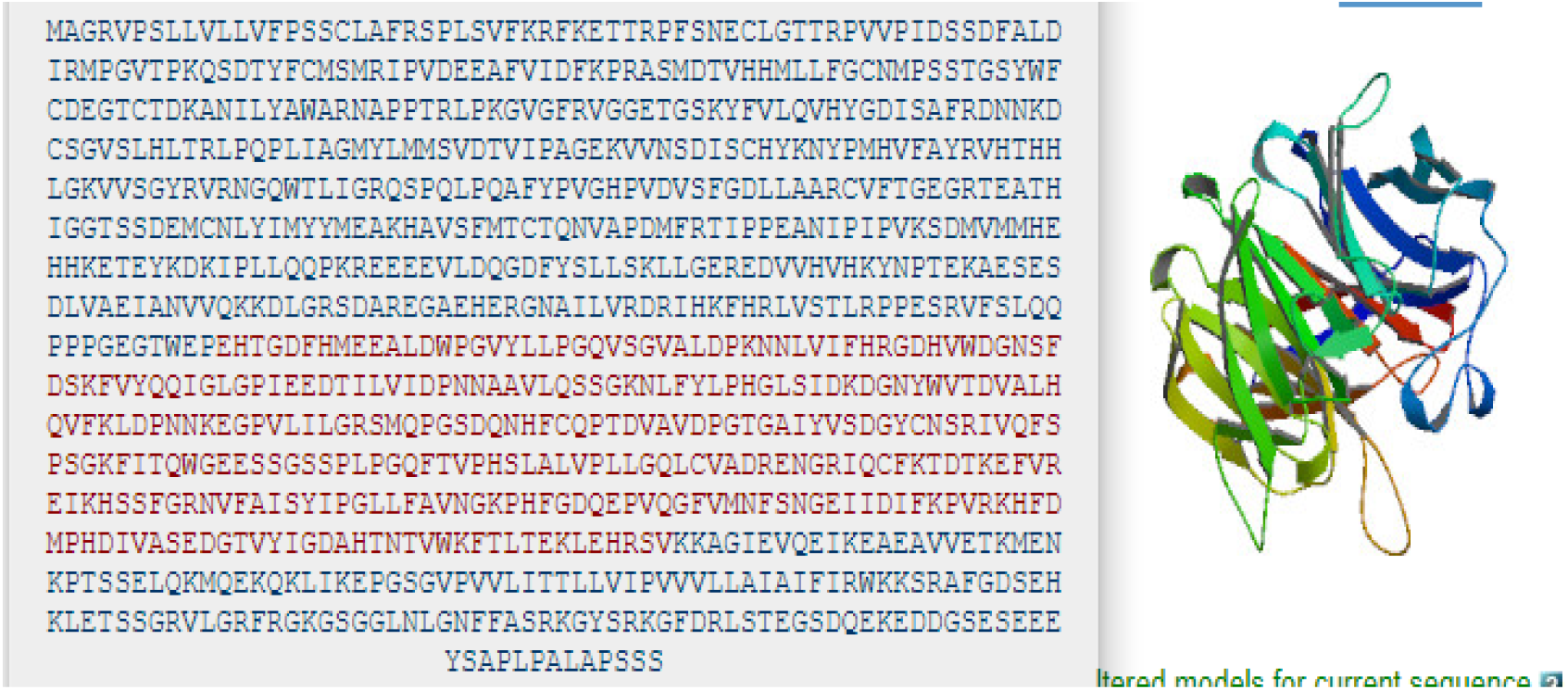
3D structure of PAM protein from Modbase.

**Figure 8.**
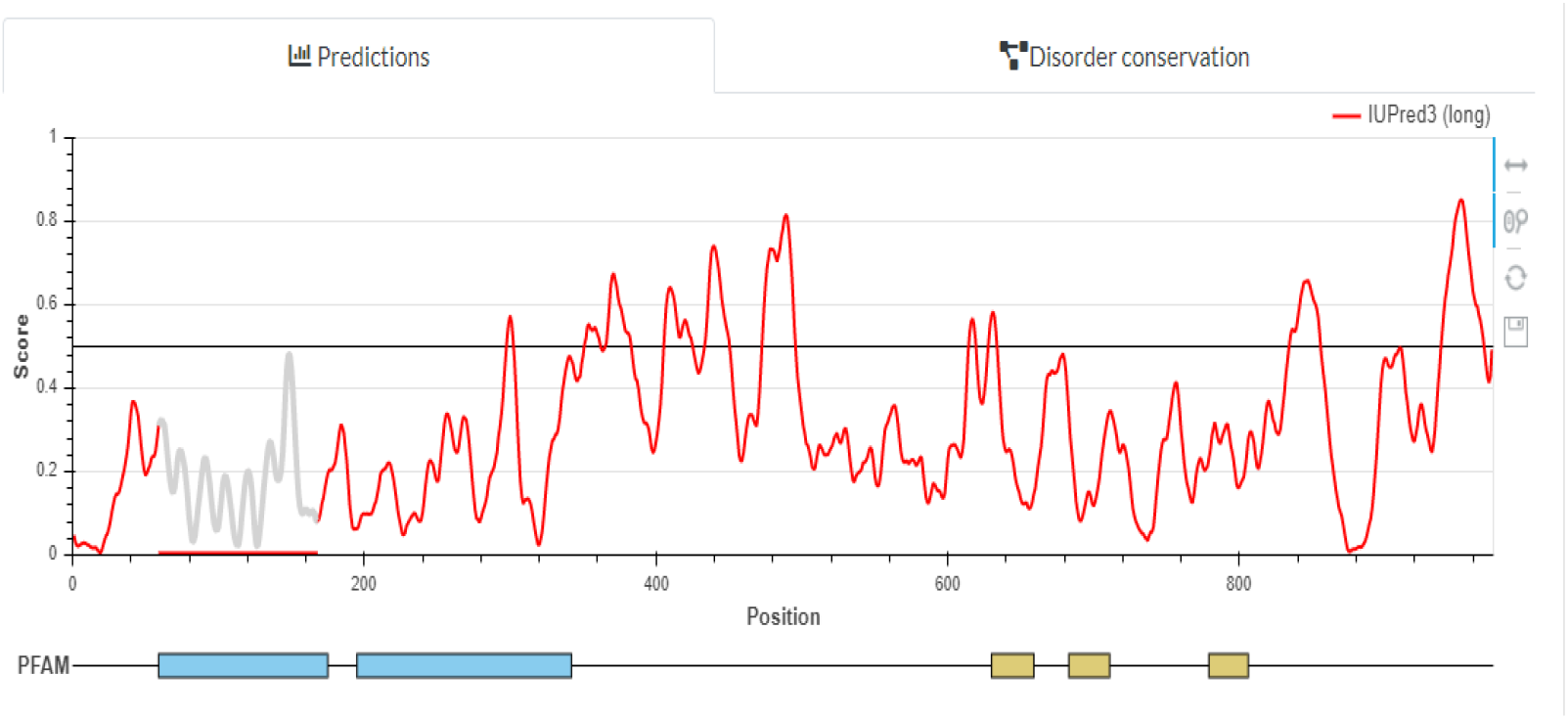
IUPred-3 disorder prediction profile.

**Figure 9.**
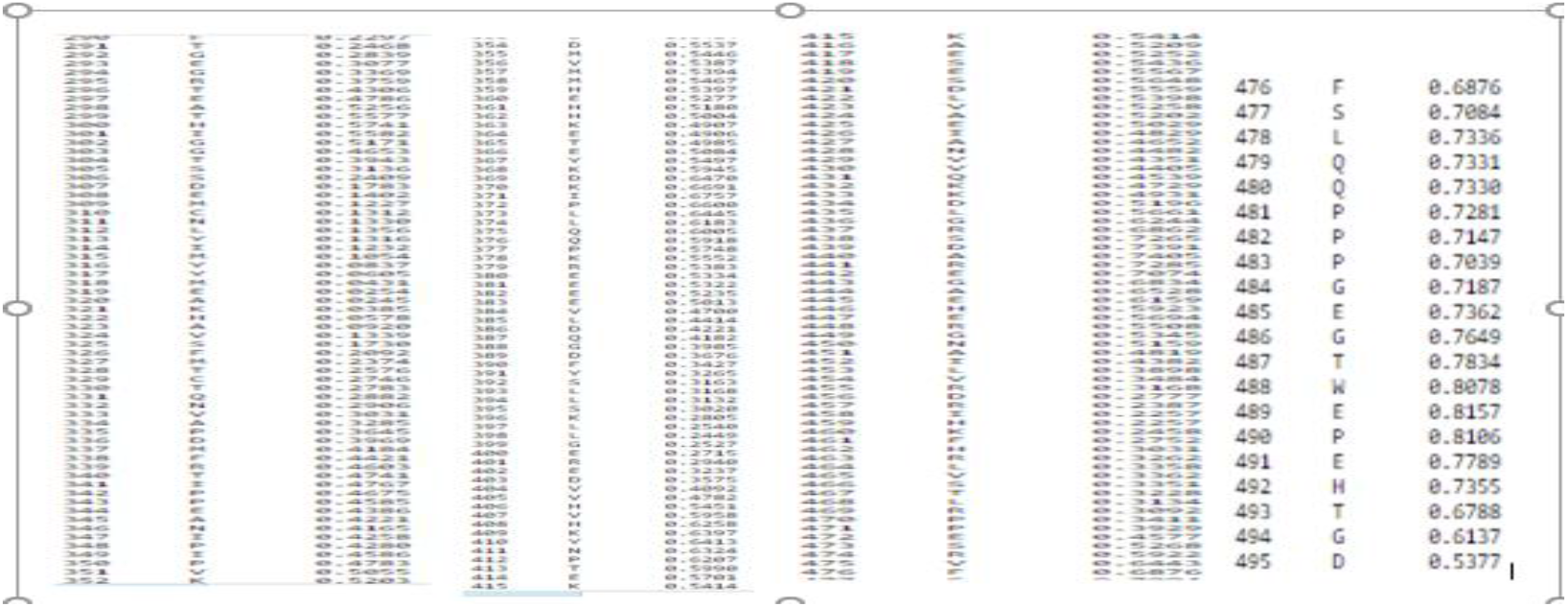
Detailed disorder scores for amino acid regions 290-495.

### Evolutionary Conservation Analysis

Comparative sequence analysis demonstrated high evolutionary conservation of PAM across species. Ensembl orthologue mapping revealed 208 orthologues with 185 showing 1:1 orthology relationship. The analysis showed highest conservation among placental mammals (108 species) and primates (26 species). BLAST analysis revealed 90% sequence similarity with Mus musculus and 89% with Rattus norvegicus. The conserved regions predominantly corresponded to catalytic domains and metal-binding sites.

**Figure 10.**
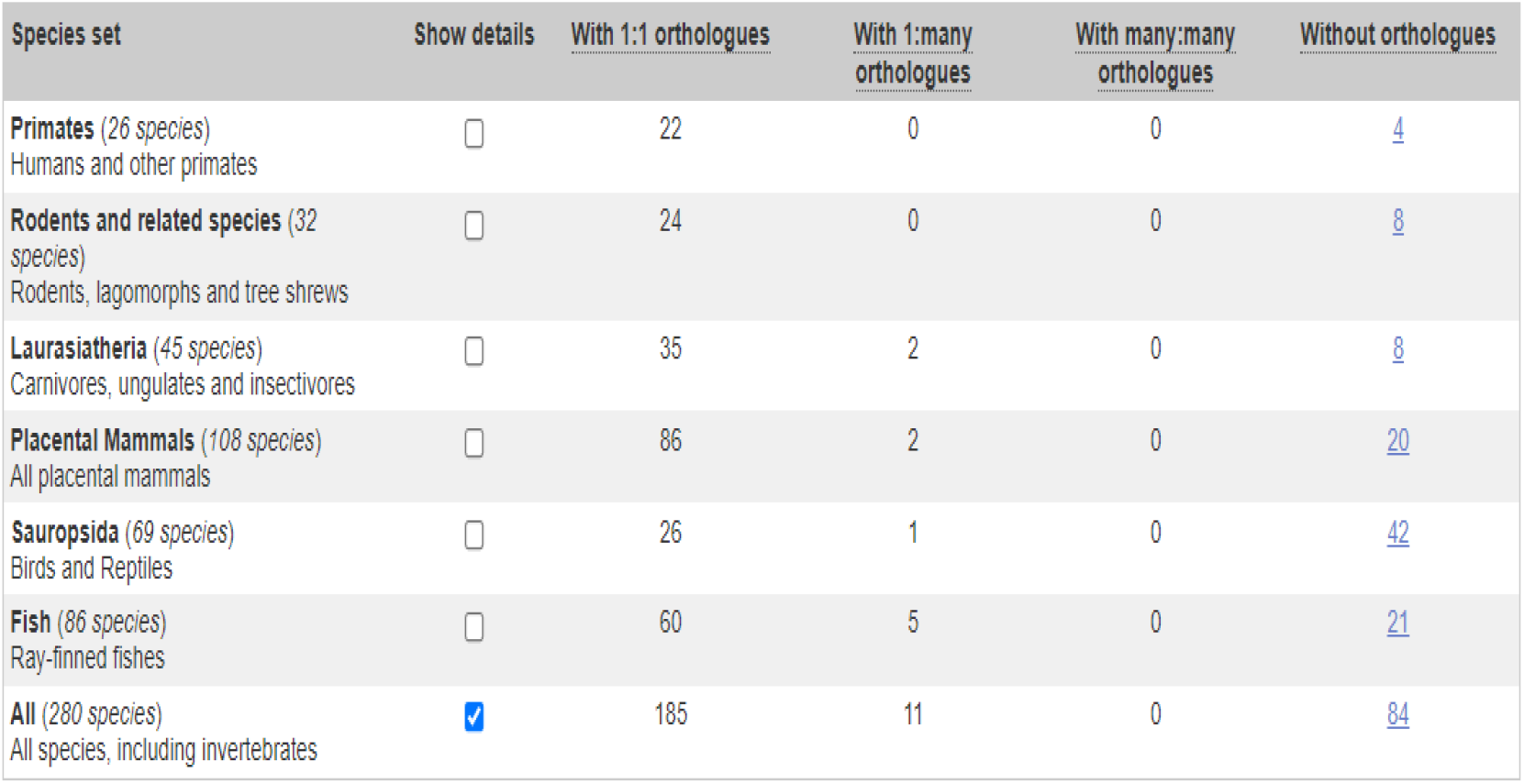
Orthologue distribution across species from Ensembl.

**Figure 11.**
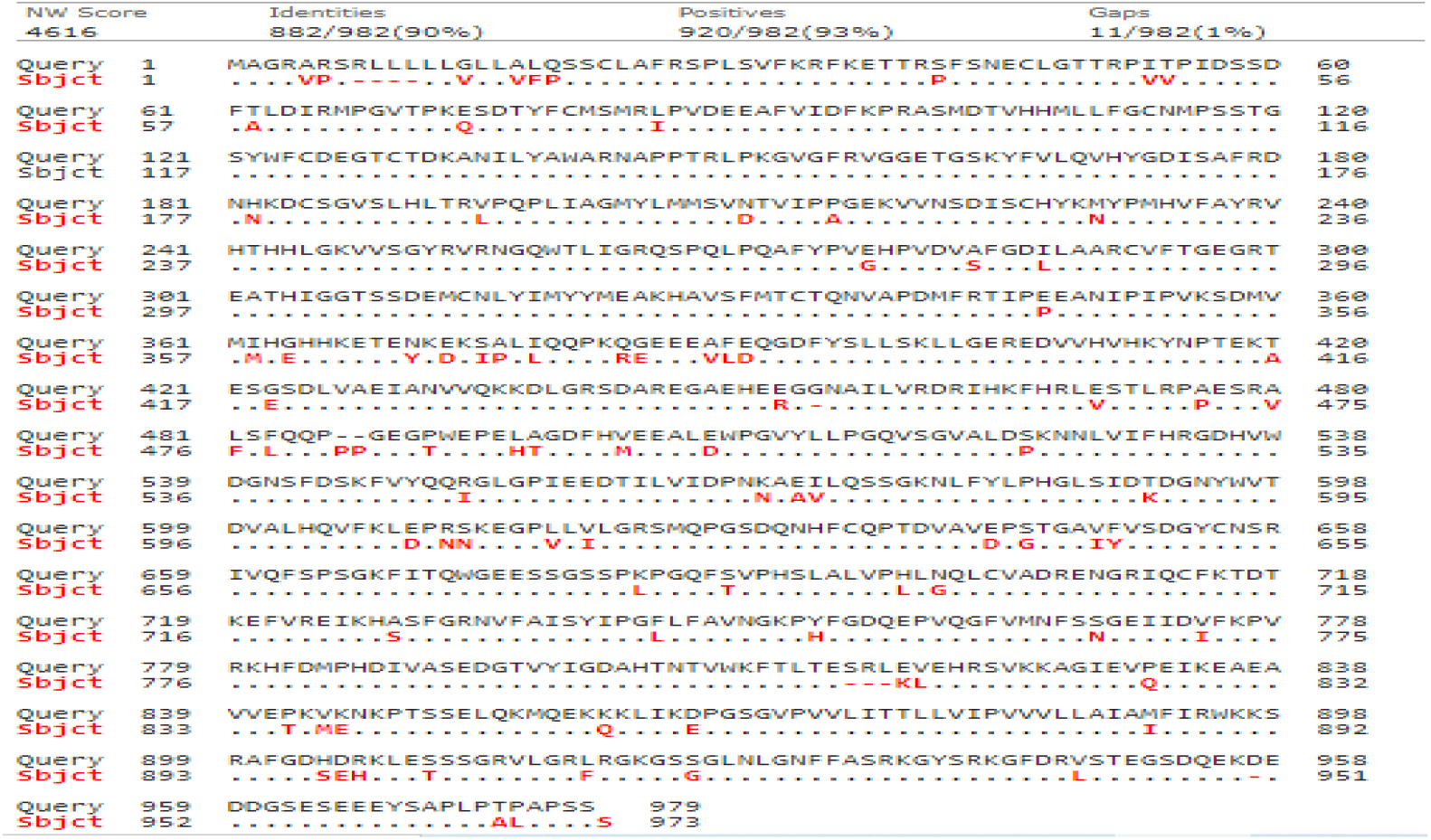
BLAST alignment with Mus musculus showing 90% similarity.

**Figure 12.**
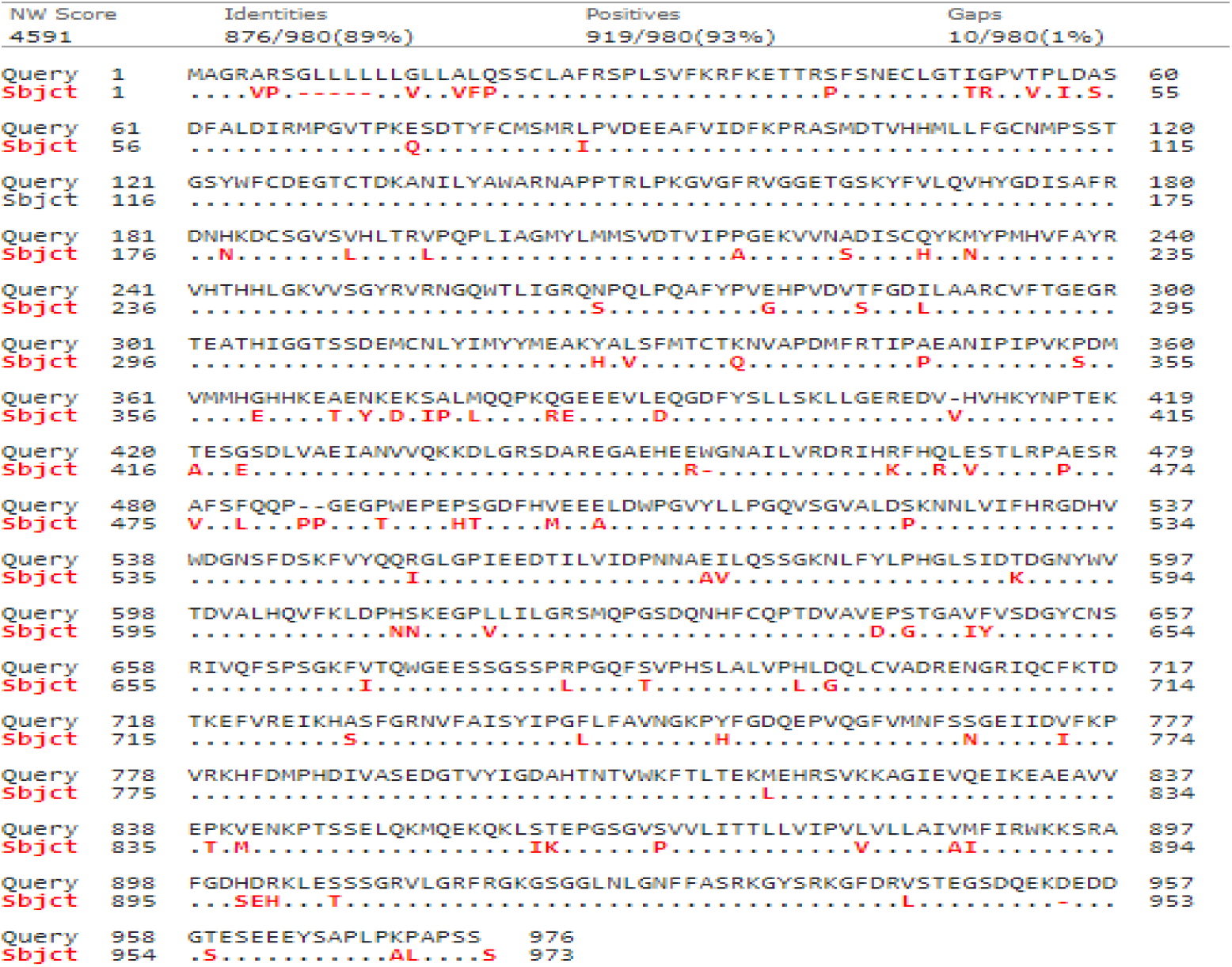
BLAST alignment with Rattus norvegicus showing 89% similarity.

### Experimental Design and CRISPR Target Identification

CHOPCHOP analysis identified specific sgRNA target sites for CRISPR-Cas9 mediated gene editing in mouse models. The tool provided rankings, target sequences, genomic locations, and off-target predictions for potential guide RNAs targeting the PAM gene locus.

**Figure 13.**
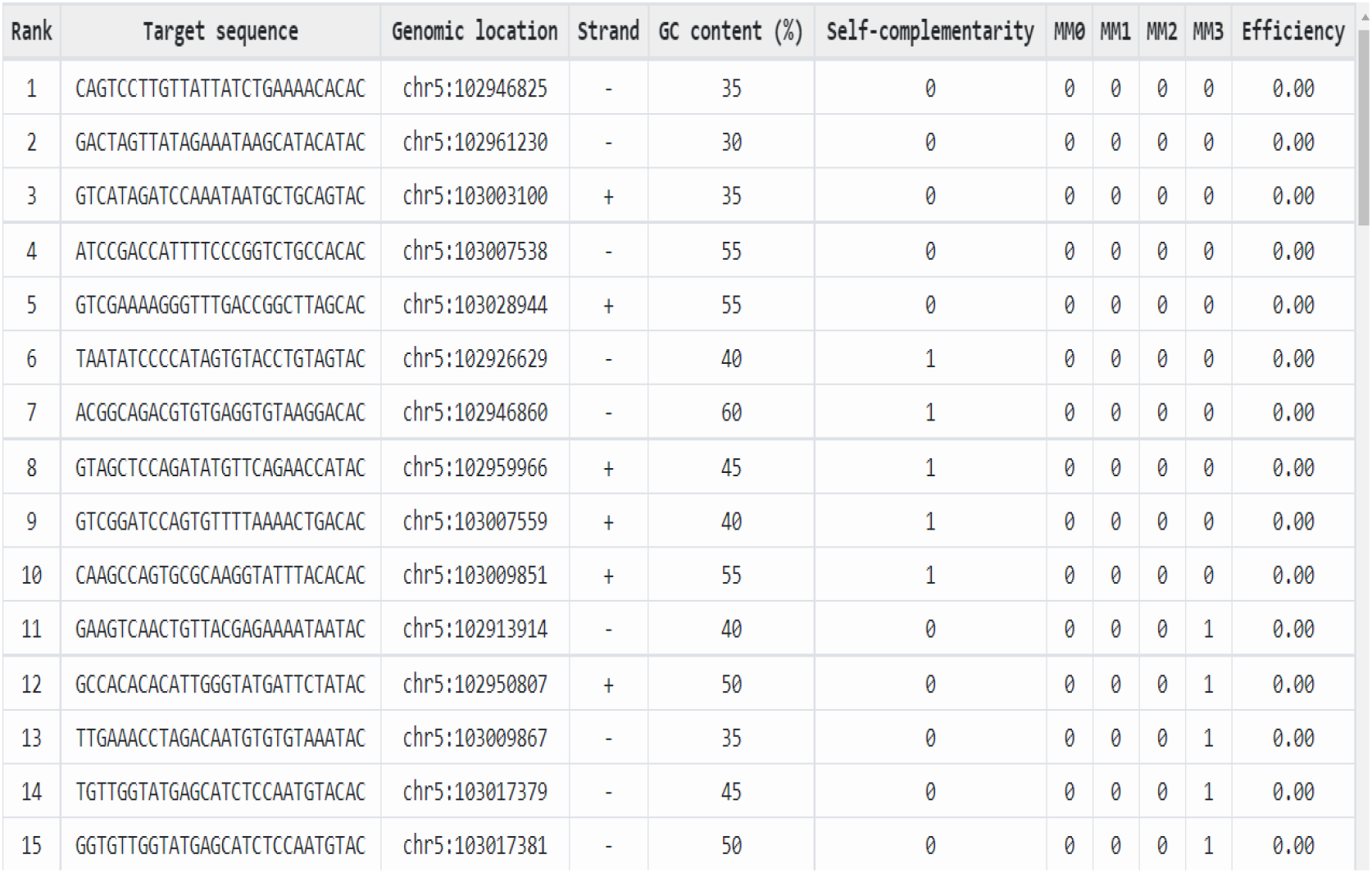
CHOPCHOP analysis for CRISPR guide RNA design.

**Figure 14.**
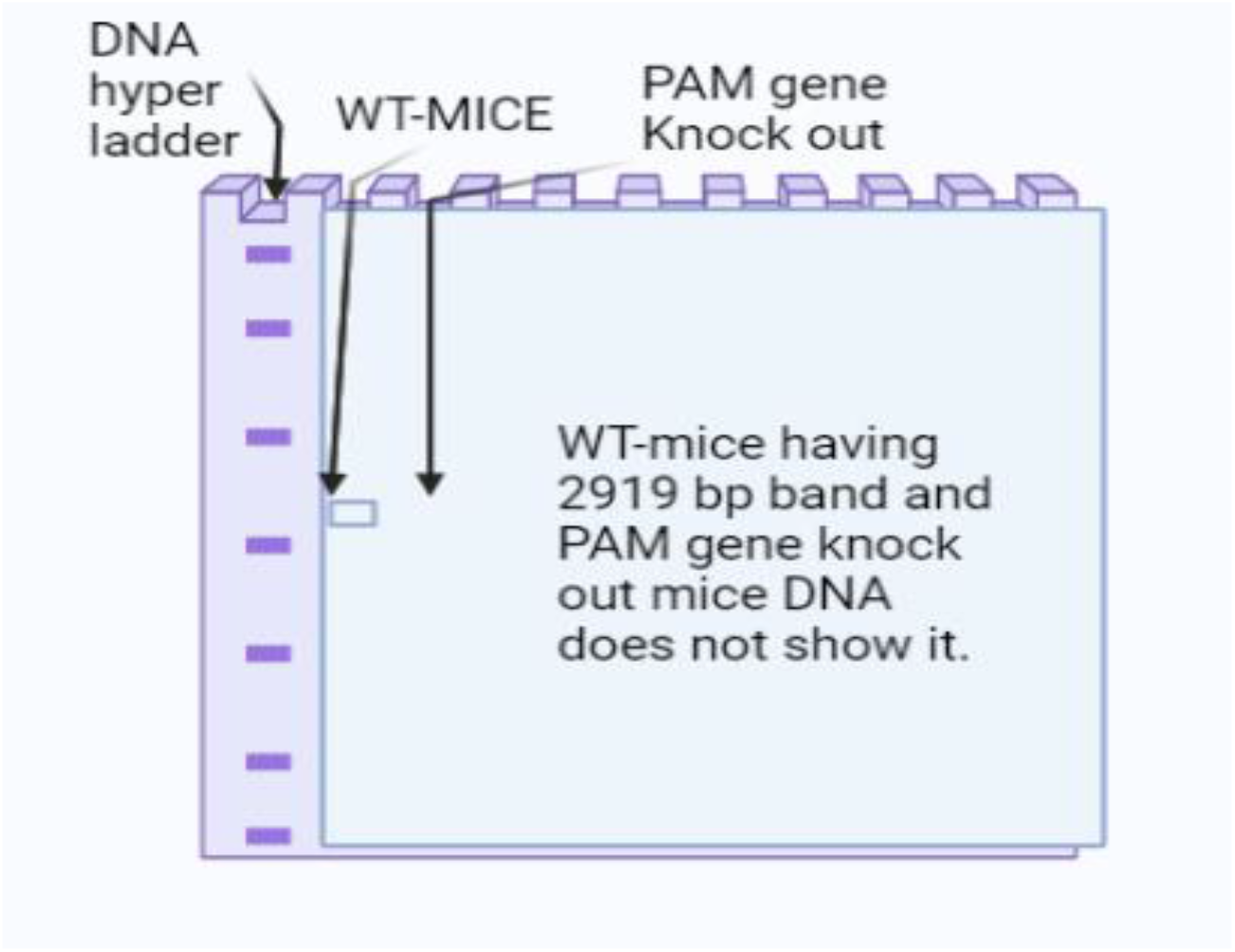
Proposed experimental workflow for PAM functional validation.

## Discussion

The comprehensive in-silico computational analysis intensively performed in this study produced detailed structural and functional data map of PAM gene. This study offers critical insights of PAM gene role in the pathogenesis of Type 2 Diabetes Mellitus (T2DM) (DeFronzo et al., 2015).

The essential catalytic domains (PHM and PAL) showed non-synonymous SNPs and highly conserved metal binding sites have prominent significance. Amidation of peptide hormones is directly impacted by abolished enzymatic activity due to impaired copper binding ability as predicted by mutations in histidine-rich regions (Eipper et al., 1993; Satani et al., 2003). This computational analyses, is strongly supported by in-vivo study elaborating the T2DM associated PAM alleles, showed reduced enzymatic catalytic activity which directly distort insulin secretion from human pancreatic β-cells (Thomsen et al., 2018).

Furthermore, the PAM gene role in protein-protein interactions is suggested by the prediction of wide intrinsically disordered region that have potential to play pivotal part in hormonal secretory pathways of islets cells through facilitating binding events. These kind of facilitations are linked with key glucose regulating hormones such as insulin and glucose that have capacity to influence the packing, processing and release of these secretory hormones (Franklin et al., 2005; Wilcox, 2005).

Common models organisms depicted high degree of evolutionary conservation. Hypotheses generated in this analyses, provide compelling validation to perform in-vivo models test (Chen et al., 2020; de Moura e Dias et al., 2021). This study identified CRISPR-Cas9 target sites offering validated and direct way for functional gene knockout models. These models can be studied to elucidate the specific SNP mutation effect on Islets cells function and modulations in homeostasis of blood glucose levels (Nagy & Einwallner, 2018).

The limitation of this study is reliance on computational predictions and analyses which set future direction for wet-lab experiments. Enzymatic activity and protein expression assays are required to confirm non-synonymous SNPs functional impact. Beside of this limitation, this multifaceted and robust in-silico framework explored here conveys meticulously high priority set of probability candidate variants and practically possible hypotheses. The work is significant as this bridge between population genetics and molecular mechanism by providing actionable and essential roadmap for future experimental studies to establish PAM gene causal role in T2DM pathophysiology and its importance as novel therapeutic target.

## Conclusion

In conclusion, this systematic bioinformatics analysis explored that impaired peptide amidation is caused by PAM gene high risk genetic splice variants and key structural elements that contribute to T2DM pathophysiology.

This work produce strong and testable hypothesis: coding variants in PAM gene interrupt its enzymatic function that leads to defective amidation of pivotal Islets peptides like glucagon and GLP-1. These peptides are involved in secretion of insulin and glucagon.

The functional domains and splice variants explored in this study should be prioritized for future insights into functional studies in cellular and animal models. The study have promising potential for PAM gene role in T2DM to assess novel therapeutic targets.

## Author Contributions

M.M. conceived the study, performed all analyses, and wrote the manuscript.

## Data Availability

All data generated or analysed during this study are included in this published article. The underlying data are publicly available from the databases cited in the Methods section (e.g., Ensembl, dbSNP, UniProt).

